# Toward Atomistic Models of Intact SARS-CoV-2 via Martini Coarse-Grained Molecular Dynamics Simulations

**DOI:** 10.1101/2022.01.31.478415

**Authors:** Dali Wang, Jiaxuan Li, Lei Wang, Yipeng Cao, Sai Li, Chen Song

## Abstract

The causative pathogen of Coronavirus disease 2019 (COVID-19), severe acute respiratory syndrome coronavirus 2 (SARS-CoV-2), is an enveloped virus assembled by a lipid envelope and multiple structural proteins. In this study, by integrating experimental data, structural modeling, and coarse-grained molecular dynamics simulations, we constructed multiscale models of SARS-CoV-2. Our 500-ns coarse-grained simulation of the intact virion allowed us to investigate the dynamic behavior of the membrane-embedded proteins and the surrounding lipid molecules *in situ*. Our results indicated that the membrane-embedded proteins are highly dynamic, and certain types of lipids exhibit various binding preferences to specific sites of the membrane-embedded proteins. The equilibrated virion model was transformed into atomic resolution, which provided a 3D structure for scientific demonstration and can serve as a framework for future exascale all-atom MD simulations.

## 1 Introduction

The ongoing Coronavirus disease 2019 (COVID-19) pandemic has infected a massive amount of people globally in the past few years. The causative pathogen, severe acute respiratory syndrome coronavirus 2 (SARS-CoV-2), is an enveloped virus assembled by a lipid envelope, a positive-sense single-stranded RNA, and four structural proteins: the spike (S), membrane (M), envelope (E), and nucleocapsid (N) proteins. For the purpose of understanding the molecular basis of viral functions, assembly, virus-host interactions, and antibody neutralization, extensive studies have been carried out to solve the structures *in vitro* for the SARS-CoV-2 viral proteins by cryoelectron microscopy (cryo-EM) or crystallography, and structures *in situ* for the native proteins by cryo-electron tomography (cryo-ET). Although recent technical developments of cryo-ET have enabled the reconstruction of intact SARS-CoV-2, the structure has been limited to nanometer resolution [1, 2].

In the meanwhile, computational studies have also provided highly valuable information on the structure and dynamics of the virus [3–7], especially the pioneering work by the Voth lab [3] and the ground-breaking AI-enabled multi-scale simulations by a large team led by the Amaro lab [4, 6]. However, the existing structural models of the virus have been either limited to a coarsegrained (CG) scale, focusing primarily on the virus envelope, or constructed without considering the specific protein localization from the *in situ* cryo-ET data, particularly the N proteins. Therefore, we set out to construct both coarse-grained (CG) and atomistic models of SARS-CoV-2 that are as intact as possible, by fully employing the latest cryo-ET data [1], the available experimentally resolved protein structures, structure prediction and modeling methods, as well as coarse-grained molecular dynamics (MD) simulations. The CG and atomistic models can not only provide 3D structures for scientific demonstration, but also offer a framework for future exascale MD simulations to understand the dynamics of the intact virus, its assembly, and its mutations.

To obtain a better equilibrated model of SARS-CoV-2, we first built a CG model and then equilibrated it with the Martini force field [8], followed by a resolution transformation. To build the CG model, we constructed structural models of each protein component separately. Then we assembled them onto a pre-equilibrated lipid envelope according to the architecture of the intact virus revealed by cryo-ET [1]. Since there is currently no way to solve or predict the full-length RNA structures within the envelope, only the N-bound RNA segments were considered in our model. The CG SARS-CoV-2 model was solved in a water box, equilibrated by a 500-ns CG MD simulation, and then transformed into the atomic resolution. Two atomistic models of the intact virus are provided: the initial structure, built according to the cryo-ET map prior to the CG MD simulation (Fig. 1a & S1c), and the final structure, built after a 500-ns CG MD simulation (Fig. S1d). Although the CG simulation was not able to quantitatively characterize the conformational changes of the viral proteins, we can obtain key information regarding the dynamic properties of the structural proteins on the envelope, such as the interactions between the membrane-embedded proteins, the diffusion coefficients of the membrane-embedded proteins, as well as the lipid clustering around them. Therefore, the CG simulation not only efficiently equilibrated the virion model for the following resolution transformation toward an atomistic model, but also provided valuable insights on the protein-membrane interaction of an intact SARS-CoV-2 virion as well as the overall dynamics of each viral component on the envelope.

**Figure 1:**
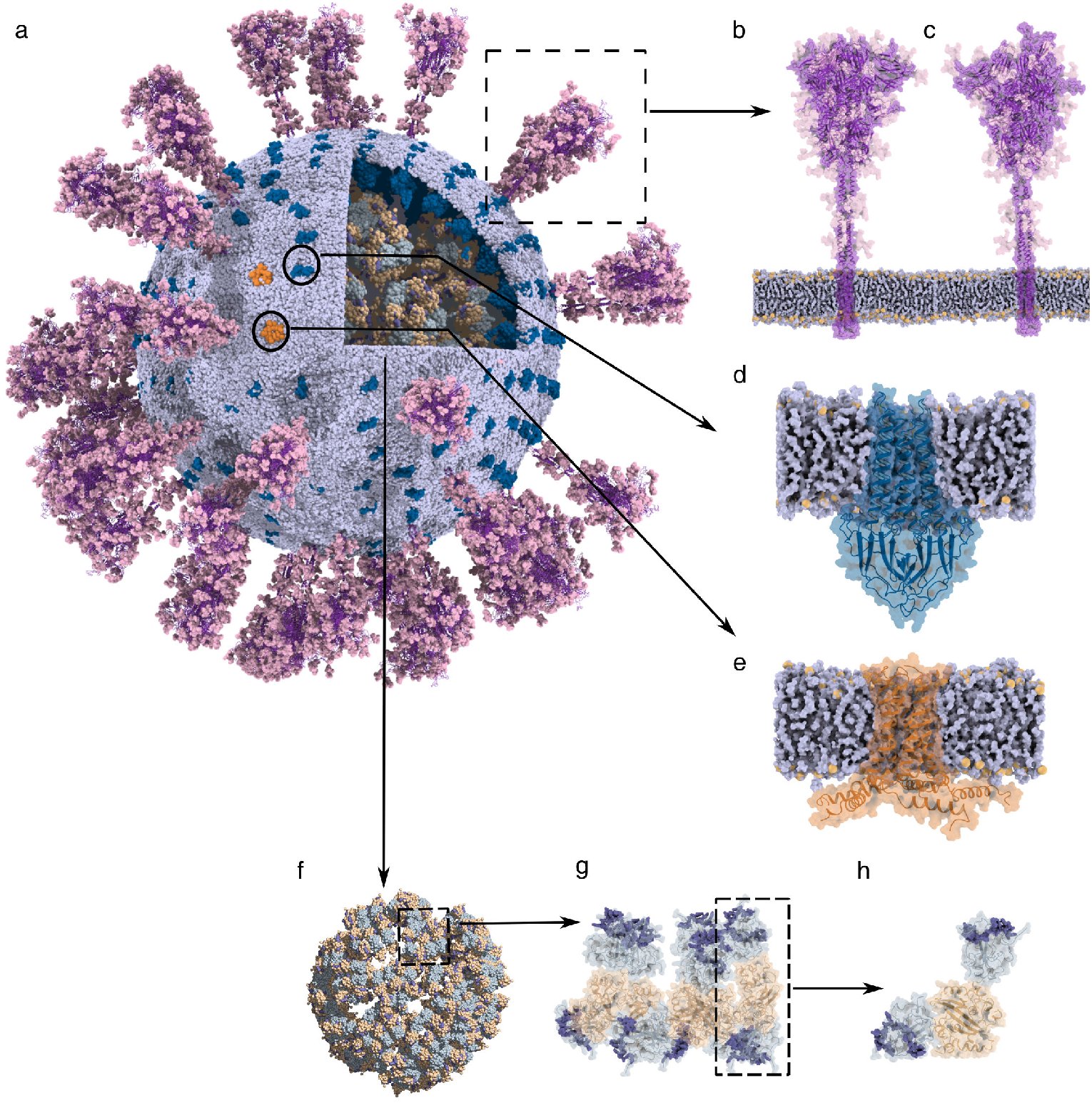
(a) The overview of the virus structure. The viral envelope is colored blue–white. Purple, deep blue, and orange regions indicate the S, M, and E proteins, respectively. RNPs are located within the envelope, and domains near the N-terminal and C-terminal of N proteins are shown in grey–blue and wheat, respectively. (b)–(c) The ‘RBD down’ and ‘one RBD up’ conformations of the S protein. The S proteins are purple, and the light-pink surface shows the glycans. The blue–white surface represents the viral envelope where S proteins are embedded, and the orange spheres indicate the lipid head groups. (d)–(e) The zoom-in view of the M protein (d) and E protein (e). (f)–(h) The architecture of RNPs: arrangement of all the RNPs within the virus (f), a single RNP unit (g), and an N protein dimer (h). The RNA segments bound to the N protein are blue–purple.

## 2 Results

Based on the cryo-ET map (EMD-30430) [1], we constructed a vesicle with a rough diameter of 85 nm as the viral envelope and assembled a virion model (Fig. 2a). Using the Martini Force Field, we were able to obtain a well-equilibrated system, in which the vesicle reached a converged size within 200 ns in the CG simulation (Fig. 2b). In the previous work, Yao et al. analyzed the SARS-CoV-2 envelope size based on 2,294 virions and demonstrated that the viral envelope shape is ellipsoidal [1]. The virion diameter measured from our CG trajectory differ slightly from the statistical data, but overall matched well with the cryo-ET map (EMD-30430).

**Figure 2:**
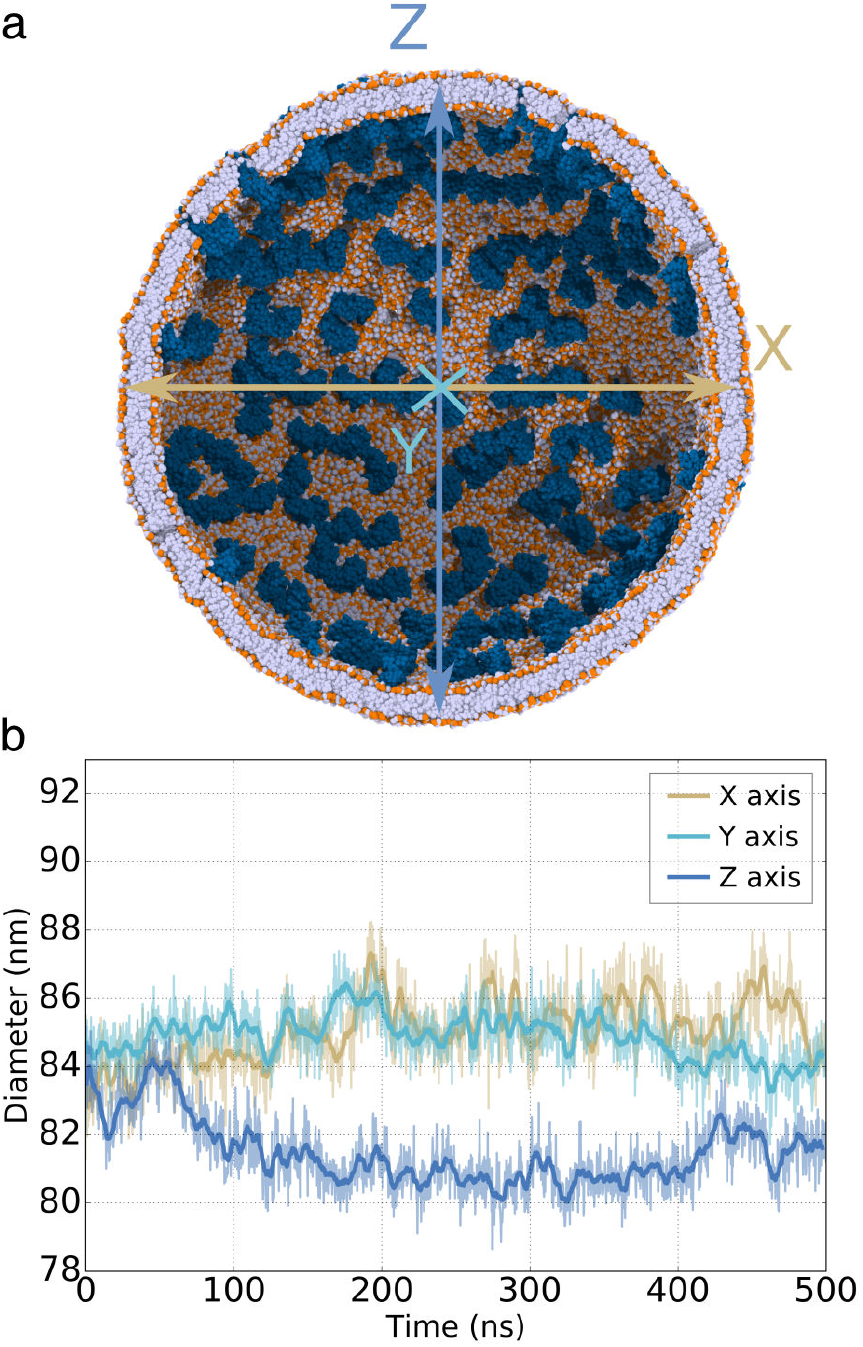
(a) The sectional view of the viral vesicle. The vesicle boundary was marked by the lipid head groups, which were colored in orange. The deep blue spheres represent the M proteins. The RNP and S proteins were hidden for clarity view. (b) Vesicle size evolution along the X (brown), Y (palegreen), and Z (lightblue) axis during the simulation. The colors of the three curves are corresponding to (a).

The protein-membrane interaction is critical along the whole life cycle of SARS-CoV-2. During the assembly stage of the virus in host cells, the membrane-bound E, M and S proteins on the ERGIC (ER-golgi intermediate compartment) recruit the viral RNPs, together budding into the ERGIC and forming new virions [9]. After the structural proteins insertion, the lipid molecules of the envelope may rearrange to find their favorite positions and cluster around the membrane-embedded proteins. We analyzed the protein-lipid interface on the virion envelope to identify the specific lipid-binding sites. We calculated the radial distribution function (RDF) of lipids around the membrane-embedded proteins. Our analysis showed that each lipid component had a similar RDF profile at the very beginning of the simulation, representing the randomly distributed lipid molecules before equilibration (top panels of Fig. 3b-d). Along with the CG simulation, the RDF profiles of various lipid molecules gradually changed and converged. As shown in Fig. 3, the RDFs calculated from the 300-400 ns trajectory (middle panels in Fig. 3b-d) and 400-500 ns trajectory (bottom panels in Fig. 3b-d) were already indistinguishable, indicating that the lipid distribution had well converged.

**Figure 3:**
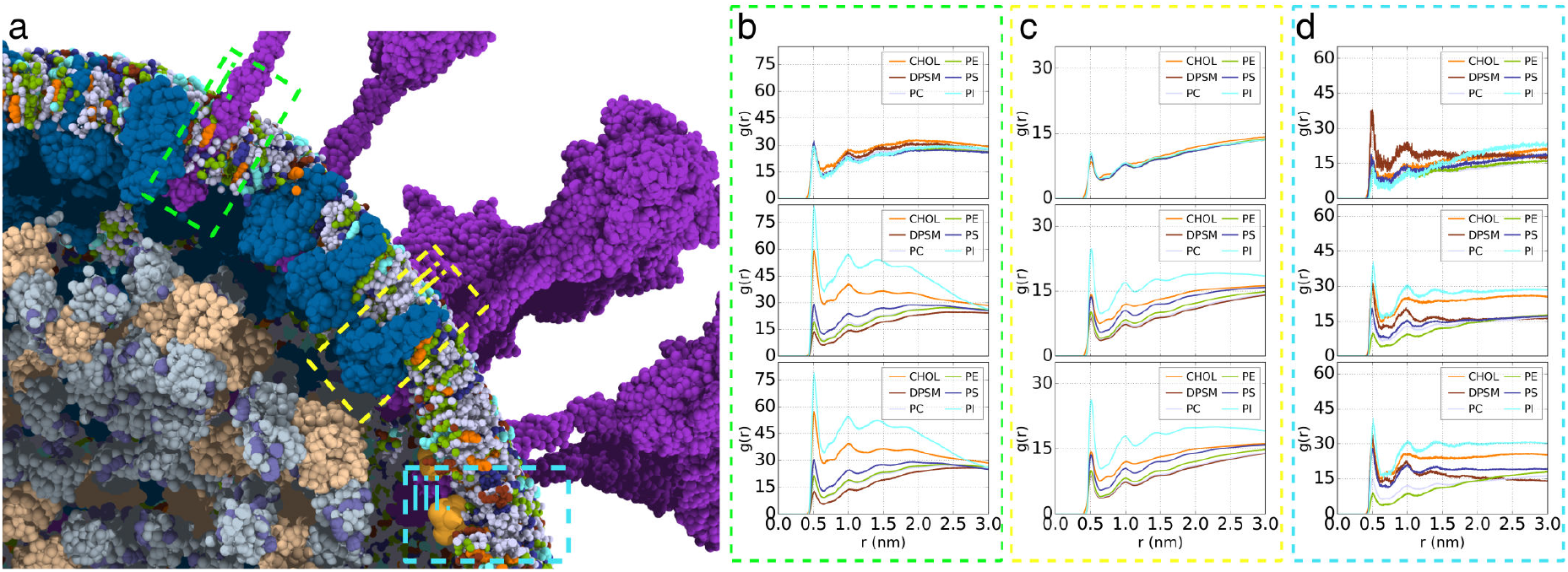
(a) A zoom view reflects the relative position between the S, M, and E trans-membrane domains and the vesicle. The colors of proteins are matched with Fig. 1. The three boxes point out the S (i), M (ii), and E (iii) trans-membrane domain, respectively. (b) The lipid radial distribution function (RDF) refers to the S trans-membrane domain. The panels from top to bottom show the RDF results generated from the 0-5 ns, 300-400 ns, and 400-500 ns trajectories. The different vesicle components were colored in orange (CHOL), chocolate (DPSM), bluewhite (PC), splitpea (PE), deepblue (PS), cyan (PI). (c)-(d) same as (b) but for M and E.

Although the transmembrane domains (TMDs) are very different among the M, S, and E proteins, the converged lipid distribution around them showed common features. Relatively, Phosphatidylinositol (PI) and Phosphatidylserine (PS) were more frequently detected around the proteins than phosphatidylcholine (PC) and phosphatidylethanolamine (PE) (Fig. 3), which illustrated that the negatively charged lipids have a stronger binding preference toward the membrane proteins in SARS-CoV-2. It was also noticeable that more PI molecules enrich around the proteins than PS, although they carry the same charge of −1 e. Further analyses showed the PI-enriched region distributes many aromatic residues (Fig. 4), illustrating that the Pi-Pi stack interaction between PI head region and aromatic residue side chain probably plays a critical role in PI-protein binding. In addition, cholesterol (CHOL) showed the second-high probability of surrounding the membrane-embedded proteins (orange lines in Fig. 3b-d). These results are consistent with previous work showing that PI and CHOL prefer concentrating around the membrane proteins [10], and a recent work by Wang et al. also reported that PI and CHOL have tendency to locate near the viral S, M, and E proteins [7]. Therefore, PI, CHOL, and PS molecules constitute the preferred surrounding environment of the TMDs of membrane proteins. The stable protein-lipid interaction interface benefits to the embedding of the structural proteins in the viral envelope and maintains the entire virus architecture during the virus life cycle. Taken together, these results indicate that the massive membrane protein insertion would significantly influence the lipid’s distribution on the virion surface, eventually forming a highly heterogeneous distribution on the stable virion envelope.

**Figure 4:**
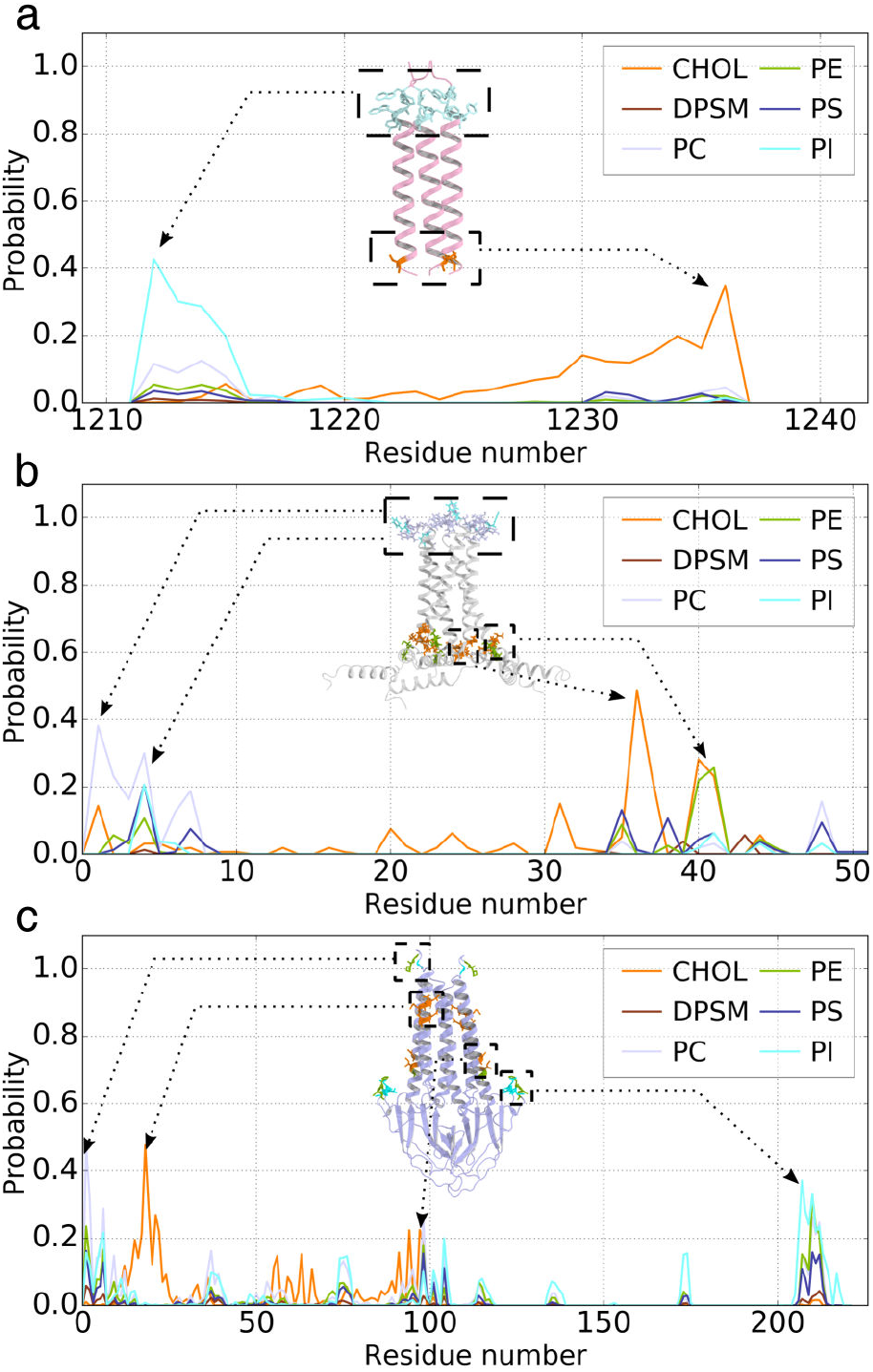
(a) Each profile represents the contact probability between S protein TMD and a kind of lipid. Various colors were applied to distinguish these profiles: orange (CHOL), chocolate (DPSM), bluewhite (PC), splitpea (PE), deepblue (PS), cyan (PI). The ribbon cartoon shows the S protein TMD structure, which is colored in lightpink. The stick representation highlights the residues with a high probability of binding to the lipid. The residues are colored corresponding to the binding lipid. (b)–(c) same as (a) but for E and M.

With the CG MD trajectory, we further analyzed how the lipids distribute around each residue of the membrane proteins to identify the specific binding sites. We calculated the lipid contact probability of each residue, and the results are shown in Fig. 4. For the S proteins (Fig. 4a), the residues in the inner and outer leaflets prefer different lipid neighbors. The CHOLs tend to locate near the inner side (M1233, L1234, C1235, C1236), while the PI lipids distribute more densely on the outer side, around W1212, P1213, W1214, and Y1215. Interestingly, we also detected that a few PC molecules were gathering near the W1212 and W1214, indicating that PC and PI may share these common binding sites. The same phenomena were observed around the E proteins (Fig. 4b), for which PI and PC lipid concentrate in the outer leaflet around S4, F6, E7, E8, while the inner-leaflet residues A36, C40, and A41 attract CHOLs. As for the lipid distribution around the M proteins (Fig. 4c), we did not observe the same asymmetric distribution of PI and CHOL as for the S and E proteins above. The PI binding sites (G6, L206, N207, T208, D209) and CHOL binding sites (L17, E18, Q19, N21, L22, S94, I97) distribute on both sides of the M proteins. However, as our structural model of the M protein dimer was mispredicted, this analysis should be interpreted with caution.

The diffusion of the membrane-embedded proteins on the virus envelope is also of high interest, which reflects how dynamic each protein is, in addition to their internal flexibility. To obtain the diffusion coefficients of M and S proteins, we analyzed the motion of each protein by calculating the mean squared deviation of all the M and S proteins in the CG trajectory (Fig. 5). The spherical coordinates *θ* and *φ* were used to describe the position of viral proteins, as shown in Fig. 5a. The motion of each protein’ scenter of geometry is shown in Fig. 5b & c, based on which we calculated the diffusion coefficient of S and M proteins to characterize their diffusion abilities (Fig. 5d & e). Our analyses showed that the M and S proteins share similar diffusion coefficients: 7.1 ± 0.2 *μm*^2^/*s* for M proteins, and 8.2 ± 1.1 *μm*^2^/*s* for S proteins, respectively. These values are close to previous CG MD simulation results, which demonstrates that the membrane protein’s diffusion coefficient in the MARTINI force field ranges from 3.3 to 12.0 *μm*^2^*/s* [11–13].

**Figure 5:**
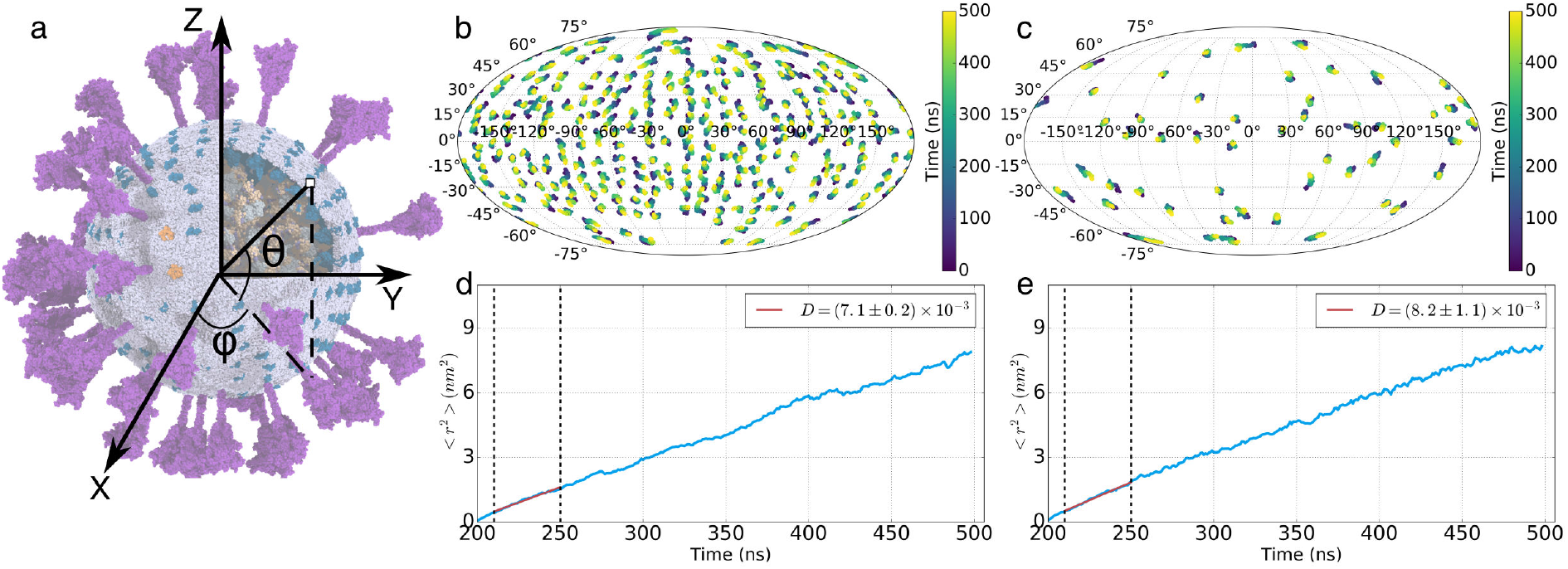
(a) Illustration of *θ* and *φ* angle. (b)–(c) The coordinate variation of the M (b), S (c) proteins transmembrane domain during the simulation trajectory. (d)–(e) The correlation between the mean squared position deviation of M (d), S (e) proteins transmembrane domain and simulation time. The dashed lines delimit the curves to do linear fitting. The diffusion coefficients of M, S proteins are 7.1 ± 0.2 *μm*^2^/*s*, 8.2 ± 1.1 *μm*^2^/*s*, respectively.

## 3 Discussion

With the whole virion model constructed (except for the complete RNA), a 500-ns CG simulation was performed in a water environment to relax each component to reach a more equilibrated configuration (Movie S1). Our simulation and analyses showed that the Martini CG simulations can be used to efficiently equilibrate such a complex system. It took at least 200-300 ns to equilibrate the virion system to reach a stable size and converged lipid distribution around membrane-embedded proteins. With such a CG equilibration, the transformed atomistic model would be more relaxed and require less computation time for further equilibration.

According to our RDF analysis, the PI lipids and CHOL were found to be more concentrated around the membrane-embedded proteins, which is consistent with another recent simulation study [7] and an earlier systematic analysis based on extensive simulations of membrane proteins [10]. The PS lipids also showed a moderate binding affinity to the S and M proteins, while the PC and PE lipids exhibited the least binding preference. The sphingomyelin (SM), DPSM, did not show binding preference to any membrane-embedded proteins either. Overall, the lipid distribution in the envelope is in line with the previous work by Corradi et al. [10]. In addition, our results showed that the PI lipids tend to concentrate on the outer leaflet, while CHOLs prefer to bind with proteins in the inner leaflet. The residues M1233, L1234, C1235, C1236 in the S protein, and A36, C40, and A41 in the E protein, located on the inner leaflet of the envelope, can recruit CHOLs. On the outer leaflet, the aromatic residues W1212, W1214, and Y1215 in the S protein, and F6 in the E protein, are PI attractive sites, indicating that the aromatic interactions may be one of the reasons for the enrichment of PIs around proteins.

Unlike a planar bilayer, the curvature of a spherical envelope may influence the diffusion of embedded proteins. Our analyses showed that the diffusion coefficients of M and S protein are 7.1 ± 0.2 *μm*^2^/*s* and 8.2 ± 1.1 *μm^2^/s*, respectively. These diffusion coefficients are close to the values calculated from a planar bilayer system [11–13], indicating that membrane proteins in a spherical membrane may have similar diffusion ability with that in a planar bilayer. Due to the smoother energy landscape in MARTINI force field, the protein in CG force field diffuses faster than in AA force field. Previous studies compared the diffusion coefficient of proteins and lipids in the CG and atomistic models [11, 14], which showed that proteins and lipids in CG may diffuse four to ten times faster than in AA models. Based on this estimation, the diffusion coefficient of M and S proteins in AA models are estimated to be around 1.8 ± 0.1 *μm*^2^*/s* and 2.1 ± 0.3 *μm*^2^*/s*, which are close to previously measured diffusion coefficients of membrane proteins (4 – 10 *μm*^2^ /*s*’) [15, 16]. It appears that the S proteins are rather dynamic, which can diffuse to form clusters (Fig. S2). These S-protein clusters may provide a more infectious condition for multiple spikes binding to one host cell receptor, which has been reported in previous studies [1, 17].

After the CG simulation completed, both the initial (Fig. S1a) and final (Fig. S1b) CG structures were transformed into atomistic models, as shown in Fig. S1c and Fig. S1d. Therefore, we are able to provide both the coarse-grained (CG) and atomistic model structures of SARS-CoV-2 here. Although there are some unavoidable uncertainties introduced by the prediction and modeling procedure, these models represent one of the most complete architectures of the intact SARS-CoV-2 so far, and they can serve as a framework for future improvements. For example, when more accurate protein structures are obtained, the structural models used here can be updated into the more reliable ones.

Unlike non-enveloped viruses, enveloped viruses are assembled by multiple structural proteins together with the lipid envelope. The presence of lipid bilayers in their assembly imposes significant challenges in the determination and simulation of intact enveloped viral structures [18]. This computational work has tried to efficiently tackled these challenges in heterogeneity through the development of an atomistic model of an authentic SARS-CoV-2 virion based on its low-resolution cryo-ET map and multi-scale modeling and simulations. Hopefully, the models will not only provide a foundation for future all-atom simulations of the intact virus, but also provide essential and intuitive information for the structural studies of enveloped viruses.

## 4 Materials and Methods

Our structural models of SARS-CoV-2 were based on recent structural biology studies, particularly the Cryo-ET density map of the virus [1], as well as protein structure prediction methods and molecular dynamics (MD) simulations. Constructing an atomistic model of such a large and complex system directly from scratch may produce massive bad contacts between atoms, which will cost a long time to relax and equilibrate. Therefore, we first built a coarse-grained (CG) model of the virion and equilibrated the system with the Martini force field [8,19]. Then the CG system was transformed into an atomistic model. The details of the system construction are as follows:

### 4.1 Construction of the membrane envelope

We set up the initial CG vesicle with the CHARMM-GUI Vesicle Maker [20]. Since the vesicle would shrink after equilibration [20], we extended the initial vesicle diameter (*D_init_* = 109 nm) to ensure it will reach the target diameter (*D_target_* ≈ 85 nm) after equilibration (Fig. S3) to match that observed in the Cryo-ET density map [1].

The detailed composition of the membrane envelope remains elusive. Previous MD simulation studies adopted various membranes with distinct lipid ratios to investigate the dynamics of SARS-CoV-2 spike protein embedded in a lipid bilayer. Hyeonuk et al. used a lipid bilayer composed of PC:PE:PS:SM:CHOL = 10:30:10:20:30, of which PE and CHOL are the majority [21]. Whereas Mateusz et al. (PC:PE:PI:PS:SM:CHOL = 50:20:15:5:5:5) and Casalino et al. (PC:PE:PI:PS:CHOL = 47:20:11:7:15) chose the membrane composition mimicking the lipid ratio of the ERGIC (ER-Golgi intermediate compartment) membrane, where PC and PE are predominant [22,23]. In this work, we followed the latter strategy to construct a complex vesicle with the composition PC:PE:PI:PS:SM:CHOL = 45:20:5:10:5:15 [24].

The CG vesicle system was pre-equilibrated in a water box of 130 × 130 × 130 nm^3^ with the Martini2.2 force field [8, 19]. After a 10000-step energy minimization, the system was equilibrated in the NPT (isothermal-isobaric) ensemble for 200 ns. The long-range electrostatics was calculated by the reaction-field method. The van der Waals interaction and Coulomb interaction were considered within 1.1 nm. The v-rescale method and Berendsen method were used to maintain the system temperature at 310 K and pressure at 1.0 bar, respectively [25, 26]. The pressure coupling was isotropic. The coupling time constants for both the pressure and temperature were set to 1.0 ps.

### 4.2 Structural model of the spike (S) protein

The initial atomistic structures of the ‘one RBD up’ (PDB ID: 6xm3) [27] and ‘RBD down’ (PDB ID: 6xr8) [28] conformations were downloaded from the Protein Data Bank (PDB). These two high-resolution structures contained most of the S protein architecture, yet there are still some residues missing. These missing residues can be categorized into two types, and we adopted distinct protocols to fill these residues with MODELLER [29, 30]:

1. The residues located in the edge of the S protein ectodomain are too flexible to determine their exact positions, which leads to unresolved gaps in the cryo-EM structures. Therefore, we modeled these disordered regions with loops to maintain the integrity of the S protein ectodomain.
2. The other unresolved structures are around the membrane envelope (residue number 1148–1273 of ‘one RBD up’ and 1163–1273 of ‘RBD down’), including the Heptad Repeat-2 region, the transmembrane domain (TMD), and the endodomain. These structures were modeled with MODELLER [29, 30], based on the secondary structure predictions by the web server SPIDER3 [31].

As the S protein is a homo-trimer, the C_3_ symmetry constraint was applied in the above modeling procedure. Then the atomistic structure was converted to the CG model for CG simulations.

Glycosylation of the specific sites on the S protein promotes the interaction between the virus and the host cell receptors, facilitating the fusion of the viral envelope, and the host cell membrane [32, 33]. Therefore, determining the specific glycosylation sites is important for atomistic modeling. Although glycosylation was not considered in the CG model or the CG simulation, we took it into account when constructing the atomistic models.

According to the previous experimental data [34, 35], numerous glycan types can be detected in each glycosylation site with different possibilities. Apart from the main N-glycosylation sites, few O-glycans are located on the three chains [23,35]. All the glycosylation sites taken into account are listed in Table S1. Here, we built the glycosylated residue sites with two criteria:

1. If one glycan type shows dominant probability, then this particular type was used to set up the corresponding glycosylated residue.
2. If multiple glycan types show similar possibilities at one site, we picked the top two probable glycan types in the glycosylated residue to represent the complex glycosylation state.

The topology file of the full-length S protein with all glycosylated sites was generated by the CHARMM-GUI GLYCAN MODELER [36] with the CHARMM36m force field [37]. The structure of the glycosylated full-length S proteins is shown in Fig. S4. The details of the glycan types on each glycosylated site are shown in Table S1.

### 4.3 Structural model of the membrane (M) protein

Previous studies showed that M proteins may form dimers on the virus envelope [38], so we built a dimeric structure of the M protein based on previous studies [38,39] with the docking software ZDOCK [40].

The structural topology of the M protein of SARS-CoV-2 and SARS-CoV (UniProtKB-P59596) should be identical or similar since they share high sequence similarity (about 96%). Previous studies on both proteins showed that the M protein of SARS-CoV and SARS-CoV-2 can be divided into two domains — the transmembrane (TM) domain and C-terminal domain (CTD) [41]. But the full-length structure of the SARS-CoV/SARS-CoV-2 M protein was not resolved. Therefore, we had to use structure prediction tools to build the M protein model. Multiple protein structure prediction methods/groups (trRosetta, FEIG-lab, AlphaFold2) give consistent two-domain architectures, but the specific predicted models vary.

As AlphaFold2 [42] was best scored in CASP14, we picked the monomeric structure predicted by AlphaFold2 to construct the M protein dimer (Fig. S5a). To obtain a rational dimer structure, we need to determine the dimer interface between the M protein monomers. A previous study illustrated that the TM domain of the M protein, which is comprised of three alpha-helices (residue 1–100), might be responsible for dimerization as well as for interacting with S proteins [38]. The CTD (residue 101-222) locates at the intracellular domain and may interact with other structural proteins such as N proteins, and is therefore excluded from the dimer interface. We limited the M-M binding area when using ZDOCK 3.0.2 and blocked the CTD atoms by changing their ACE type to 19 in the PDB file. Then we followed the common procedure of ZDOCK and predicted 2000 possible complexes for evaluation and selection. The most probable and reasonable dimeric model for the construction of the virus structure (Fig. S5b) was chosen under these criteria: the TM domain and CTD in the dimer maintain the same ’up’ and ’down’ orientations; the two monomers keep certain symmetry, especially the TM domains and their parallel helices; the CTD should not intrude to the membrane region nor crash with intracellular proteins such as the RNPs in the cryo-ET density map.

After our modeling and simulations completed, the dimeric structure of M protein was resolved [43]. There are differences between the predicted structure and the resolved structure, such as the relative positions between the three transmembrane helices and the tilt angle between the transmembrane domain and the CTD (Fig. S5c & d). However, the overall scaffold is similar. Not only the secondary structure of the transmembrane domain but also the CTD are coincided between the predicted and the resolved structures. In addition, the size of the transmembrane domain in the predicted structure is similar to the resolved structure, so we think the predicted M structure can be used for rough modeling and CG simulations of M proteins, which can then be replaced with the experimental structure for further simulations.

### 4.4 The envelope (E) protein

The E protein structure was published (PDB ID: 7k3g) [44], in which the transmembrane domain was resolved, whereas the N-terminal loop and endodomain structure remain uncertain. The secondary structure prediction by the RaptorX and SPIDER3 web servers [31, 45] indicates that the endodomain of E protein may form an alpha-helix, but the orientation of this inner helix cannot be determined. The homology modeling structure (Fig. S6a) based on the SARS-CoV E protein looks strange as the endodomain helices roll up toward the TMD helices, meaning that the endodomain helices are inserted into the viral envelope. However, our recently developed membrane contact probability (MCP) predictor [46] showed that while the residues 8–34 (forming the transmembrane helices in resolved E structure) entirely interact with membrane with high probability (Fig. S6b, green curve), the inner helices (residue 38–60) (Fig. S6b, blue curve) show discrete MCP signal, reflecting that the inner helix may be adsorbed onto the membrane surface rather than being embedded into the membrane. Interestingly, the E protein structure predicted by Feig’s Lab [47] showed that the endo helix is optimized to touch the viral envelope, which is consistent with our MCP prediction (Fig. S6c). Also, the E protein structure predicted by the Feig Lab has been proven to be stable in microsecond MD simulations [48]. As for the oligomerization state, previous studies showed that the E protein of coronaviruses (like MHV and SARS-CoV) is able to self-oligomerize to create a pentameric ion channel, making this protein a viroporin [9, 49]. Therefore, we picked the predicted structural model by the Feig Lab as the initial E protein structure for our virus construction.

### 4.5 The nucleocapsid (N) protein

The N protein monomer contains five domains: the N-terminal domain (NTD), RNA binding domain (RBD), central Ser/Arg (SR)-rich linker, a dimerization domain and C-terminal domain (CTD) [9]. Previous studies have reported that the critical residues responsible for RNA binding are located in the N terminal region of N proteins (NTD and RBD) in multiple coronaviruses [50–53]. The dimerization domain is thought to mediate the formation of the N protein dimer. The RBD and dimerization domain are separated by the SR-rich linker, which is an intrinsically disordered region (IDR). In addition to the SR-rich linker, the N-terminal loop and C-terminal loop of N protein are both IDRs as well [54].

The N-terminal RBD (PDB ID: 7act) [55] and C-terminal dimerization domain (PDB ID: 6yun) [56] structures of the SARS-CoV-2 N protein have been determined separately. While these two isolated structures cannot tell the interface between them. The viral ribonucleoprotein (RNP) cryo-ET density map (EMD-30429) [1] provides a paradigm about how these two domains bind, which guided us to perform protein-protein docking with ZDOCK [40] to construct an N protein dimer (Fig. S7a).

We did not fill the IDRs of the N protein in our model. Instead, we utilized distance restraint to maintain the N protein structure (details in the next section). The full-length RNA was not included in our model because there is no way to determine the whole 3D RNA structure at the moment. However, the RNA fragment (10 bps) with a definite structure resolved together with the N protein (PDB ID: 7act) was included in our model. As the recent cryo-ET density map (EMD-30429) showed, the viral RNP unit was composed of five N protein dimers. Thus, the RNP unit structure was obtained by aligning five N protein dimers into the density map (EMD-30429) (Fig. S7b).

### 4.6 Assemble of the SARS-CoV-2 virus

After the envelope and structural proteins were set up as described above, we assembled all the components into one piece according to the Cryo-ET density map (EMD-30430) that clearly identified the architecture of the entire virus [1].

Firstly, we used the ‘fitmap’ tool within Chimera [57] to fit the equilibrated vesicle and S proteins into the cryo-ET density map (EMD-30430). Since most of the S proteins ectodomain show significant tilt with respect to the normal axis of the envelope in the cryo-ET density map, rigidly aligning our S protein model structure into the corresponding density often results in inappropriate orientations of S proteins, where their transmembrane domains were not embedded in the lipid bilayer (Fig. S8a). Therefore, after the rigid alignment, we optimized the orientation of each S protein to make its first principal axis parallel to the normal axis of the envelope and moved each S protein along the membrane normal direction to embed the transmembrane domain into the viral envelope properly. After the optimization, S proteins were located at the viral surface with an initial orientation perpendicular to the membrane surface (Fig. S8b). Then the optimized S protein structures were transformed to CG model in the Martini force field [8, 19]. Usually, the elastic network (ELN) algorithm is used to maintain the global protein conformation during the CG MD simulations. A longer ELN cutoff will enlarge the ELN intensity and make the protein more rigid. From the cryo-ET data [1], it was observed that the S proteins tend to tilt 40° relative to the normal axis of the viral envelope. To reproduce this flexibility of S proteins, we performed a series of simulations with an S protein embedded into a lipid bilayer with different ELN cutoffs. From the tilt angle analysis (Fig. S9), an ELN cutoff of 0.8 nm showed the largest flexibility and reasonable orientation angles of the S protein on the lipid bilayer surface within the simulation time, which was therefore adopted in our CG MD simulations for the S proteins. Please note that the utilization of an ELN may introduce some artifacts to the dynamics of S proteins, which is an intrinsic limitation of the CG MD simulations, but this would not be an issue for the model constructing purpose at this stage.

Next, 32 RNP units (Fig. S7b) were fitted into the density map (EMD-30430), where all the RNPs are nestled up to the inner surface of the viral envelope (Fig. S10a). Then we transformed all the RNP structures into a CG model. Without full-length RNA binding, the assembled RNPs may be unstable, so we applied distance restraints (force constant was set to 1000 kJ mol^−1^ nm^−2^) between each pair of N protein dimer to maintain the relative positions of the RNP units during the following simulations (Fig. S10b). To maintain the entire RNPs architecture we also applied distance restraints (force constant 1000 kJ mol^−1^ nm^−2^) between the center of mass (COM) of each RNP unit and the COM of all the RNPs. In addition, as all the IDRs of N protein are absent in the dimer structure, which may cause the N protein structure dissociation during the CG MD simulations, we utilized ELN (cutoff = 2.0 nm) to maintain the overall stability of the N protein dimers as well (Fig. S10c).

M proteins are located in an intricate lipid environment and are hard to be distinguished from the density map (EMD-30430). Therefore, it is difficult to directly fit the M proteins into the Cryo-ET density as had been done for S proteins and RNPs. Previous studies showed that the ratio of M:N proteins ranges from 1:1 to 3:1 [38, 58]. In the Cryo-ET density map (EMD-30430), there are 32 RNPs (32 × 5 N protein dimers) per virus. Therefore, 320 M protein dimers (M:N = 2:1) were initially inserted into the viral envelope uniformly with random orientations, and then the M protein dimers orientations are adjusted to ensure that the transmembrane domains are fully inserted into the envelope, and the first principal axis of M dimer is parallel to the normal direction of the envelope. After optimizing the orientations of the M proteins, there were 66 M protein dimers showing bad contacts with other structural proteins. As this will lead to infinite energy in the following energy minimization and equilibration procedure, we removed these 66 M protein dimers with bad contacts. As a consequence, there were 254 M protein dimers left in the system, still resulting in a reasonable ratio of M:N ≈ 3:2.

Like M proteins, E proteins are also embedded into the viral envelope, whereas far fewer E proteins are detected in a mature virus, as previous studies showed that the ratio of M:E ≈ 100:1 [59]. Therefore, we replaced two M protein dimers with E protein pentamers with proper orientation.

Following the above procedure, we assembled all the structural proteins (50 Spikes, 160 N dimers, 252 M dimers, and 2 E pentamers) into the viral envelope to form a SARS-CoV-2 virus in the absence of the complete RNA. We removed the lipid molecules within 0.1 nm of the proteins and solvated the protein–vesicle system into a cubic box of water molecules. In the end, the 155 × 155 × 155 nm^3^ sized simulation box contained 31,226,794 CG beads in total.

### 4.7 Coarse-grained molecular dynamics simulations

All the MD simulations were performed with the software GROMACS 2018.4. The CG simulation system was first performed an energy minimization using the steepest descent algorithm for 30,000 steps, followed by equilibration in the NPT ensemble (constant pressure and constant temperature) for 25 ns with the time step gradually enlarged from 1 fs to 5 fs. The Berendsen algorithm was utilized to maintain the system temperature at 310 K and pressure at 1.0 bar [26]. The coupling constant Tau-T and Tau-P were set to 1.0 ps. The pressure coupling type was set to isotropic, and the compressibility was 4.5 × 10^−5^ bar^−1^. The electrostatic interactions were calculated with the reaction-field method. The van der Waals interaction was cut off at 1.1 nm. After the equilibration, we performed a 500-ns CG MD production simulation with the time step of 5 fs.

### 4.8 Trajectory analysis

#### 4.8.1 Vesicle size measurement

To evaluate the transformation of the viral envelope shape in the CG trajectory, we analyzed the vesicle diameter along the X, Y, and Z axes during the simulation. Through the center of geometry of the vesicle (COG_vesicle) and along the X axis direction, we delimited a cylinder whose radius was set to 1 nm. Then we extracted the lipid PO4 bead within the cylinder and classified these beads into two categories: one category contains the beads whose X coordinates are less than the x coordinate of the COG_vesicle, while the other category contains the rest. The distance between the COG of each category was used to character the vesicle size in the X direction. The same procedure was adopted to assess the vesicle size along the Y and Z direction.

#### 4.8.2 Radial distribution function (RDF) analysis

The RDF profiles were generated with gmx_rdf, a built-in analysis tool of GROMACS. The lipids within 5 nm of the proteins were considered into the RDF calculation. We performed RDF analysis towards every S, M, E proteins and averaged the corresponding results to represent the lipid distribution around these embedded structural proteins. In addition, to examine whether the RDF has converged we picked three trajectory segments 0-5 ns, 300-400 ns, 400-500 ns for RDF calculation.

#### 4.8.3 Protein-lipid interaction

The protein-lipid interaction was considered when the distance between protein residue and lipid head group is less than 6 Å. We counted the frames (frames_interact) that generic lipid L can interact with protein residue R in the last 100-ns trajectory. Then the ratio between the frames_interact and total frames of the trajectory (frames_total) was used to reflect the probability that residue R contacts with the lipid L. Fig. 4 showed the average results among 50 Ss, 252 Ms and 2 Es.

#### 4.8.4 Diffusion coefficient

The protein motion on the vesicle surface can be viewed as a 2-dimensional diffusion. The position of each protein can be described by two coordinates, the latitude (*θ*) and longitude (*φ*) with respect to the COG of the vesicle (Fig. 5a). In Fig. 5b & 5c, we plotted these coordinates with Mollweide projection and colored these data points by their time stamps. The displacement of a protein, *r*, can be characterized by the arc length on the sphere surface, which was utilized to further calculate the mean squared deviation (< *r*^2^ >) of all the proteins. Given the positions of a protein before and after a short interval, (*θ*_1_ *φ*_1_) and (*θ*_0_, *φ*_0_), we can calculate the displacement of the protein by:

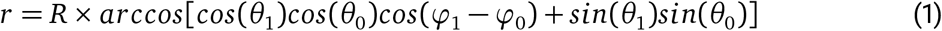

where *R* is the radius of the viral envelope.

Then the mean squared deviation (< *r*^2^ >) can be calculated, and so is the diffusion coefficient according to:

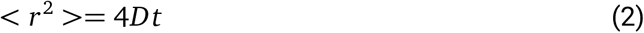

where t is the simulation time.

As our analysis showed that the system needed 200 ns to reach equilibrium in size, we only used the trajectories from 200 to 500 ns for the diffusion coefficient calculation. The data from 210 to 250 ns (delimited by dashed lines in Fig. 5d & 5e) was extracted to perform a linear fit to calculate the diffusion coefficient of proteins.

### 4.9 Conversion from the CG system to the atomistic system

Here we present two all-atom virus models transformed from the first and last frames of CG simulation by the CG2AT2 tool [60]: (1) The all-atom model of SARS-CoV-2 virion converted from the initial CG structure contains 15,526,323 atoms, in which all the S proteins are full-glycosylated. Totally, there are 278,131,974 atoms in the entire atomistic system of the simulation box. (2) Because the glycosylation did not be considered in the previous CG simulation, directly transformed the final CG virus structure to atomistic resolution results in the loss of glycosylated residues in this atomistic virus model, whose atom number is 14,873,073. Correspondingly, the simulation system involves 266,063,412 atoms after solved the virus structure into a water box. Although challenging at the moment, these atomistic systems can serve as the initial structure for future all-atom simulations.

## Author Contributions

C.S. and S.L. conceived the idea and supervised the project. D.W. and J.L. built the models and performed molecular dynamics simulations. L.W. and Y.C. participated in protein modeling. D.W., J.L., S.L., and C.S. wrote the original manuscript.

## Acknowledgements

This work was supported by grants from the National Natural Science Foundation of China (21873006 and 32071251 to C.S.; 32171195 to S.L.), the Ministry of Science and Technology of China (2021YFE0108100 to C.S.), and Tsinghua University Spring Breeze Fund (#2021Z99CFZ004 to S.L.). D.W. was supported in part by the Postdoctoral Fellowship of Peking-Tsinghua Center for Life Sciences. The MD simulations were performed on the Computing Platform of the Center for Life Sciences at Peking University and the National Supercomputer Center in Tianjin.

