## Supplementary Information for "Toward Atomistic Models of Intact SARS-CoV-2 via Martini Coarse-Grained Molecular Dynamics Simulations"

### 1 Supplementary Tables

Table S1. The glycan types of the glycosylated sites of the S protein

| | Chain A | Chain B | Chain C |
|---|---|---|---|
| N17, N149<br>N1098 | 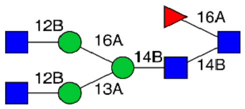 | 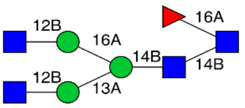 | 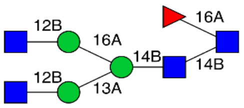 |
| N61, N603<br>N709, N717 | 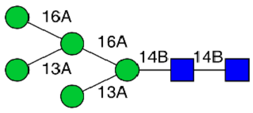 | 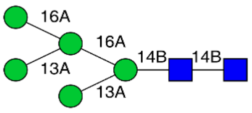 | 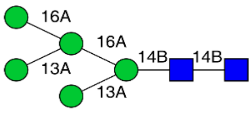 |
| N74 | 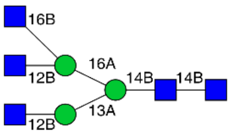 | 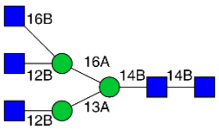 | 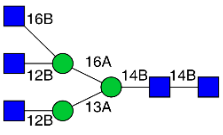 |
| N122, N801 | 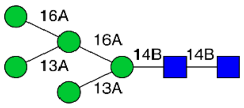 | 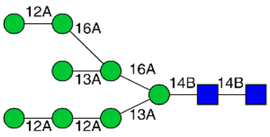 | 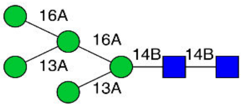 |
| N165 | 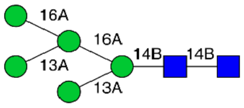 | 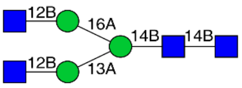 | 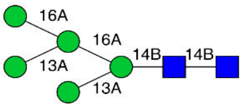 |
| N234 | 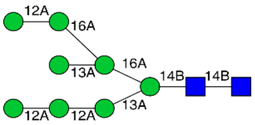 | 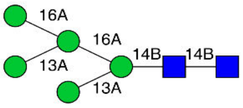 | 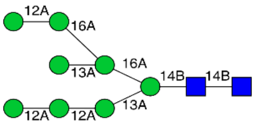 |

■ GlcNAc ● Mannose ▶ Fucose

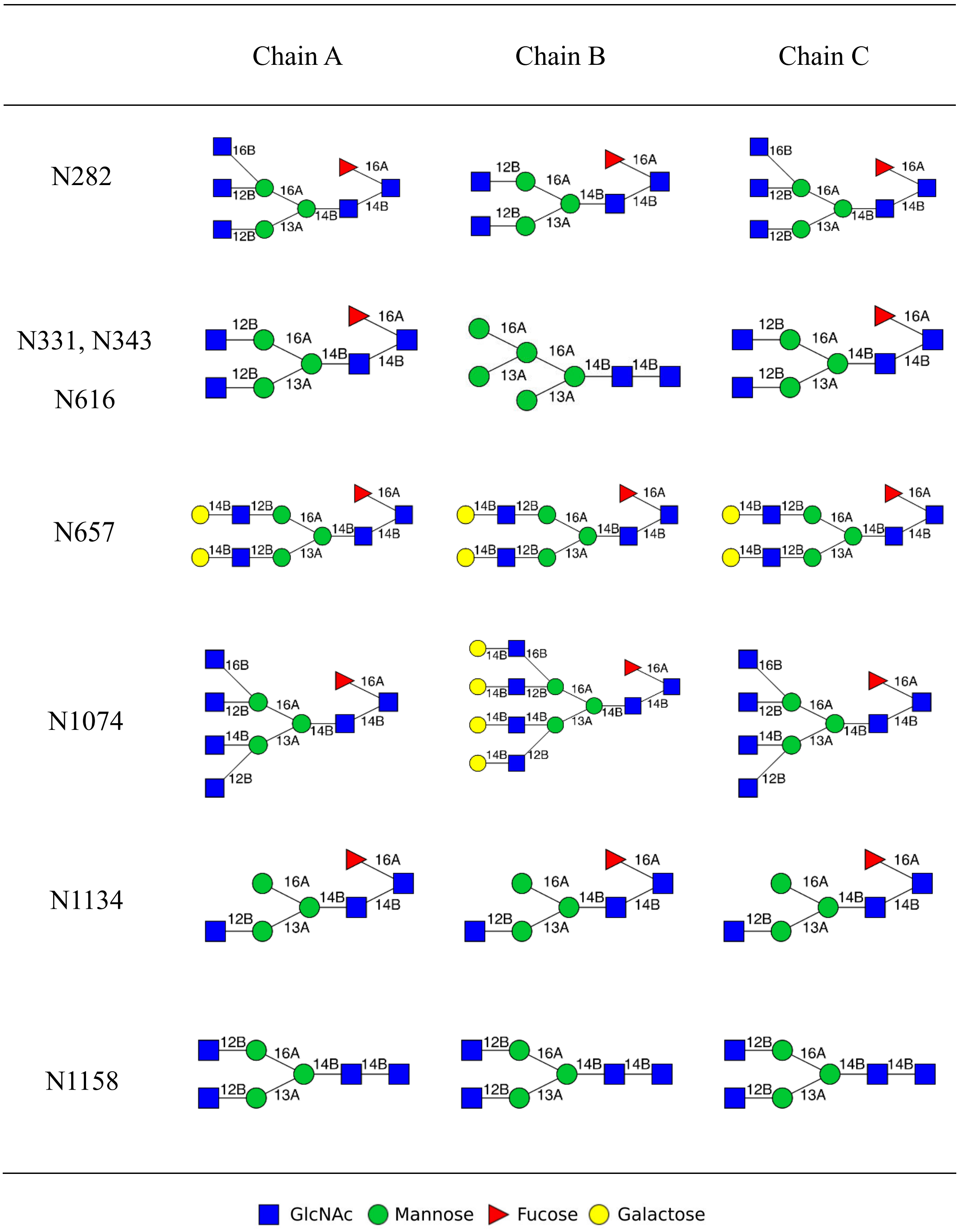
Chain A
Chain B
Chain C
N282
N331, N343
N616
N657
N1074
N1134
N1158
GlcNAc
Mannose
Fucose
Galactose

| | Chain A | Chain B | Chain C |
|---|---|---|---|
| N1173 | 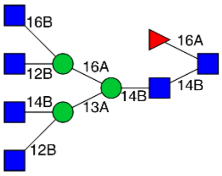 | 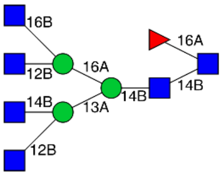 | 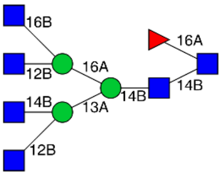 |
| N1194 | 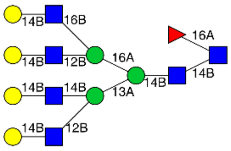 | 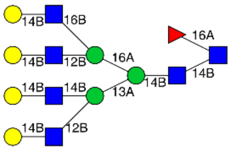 | 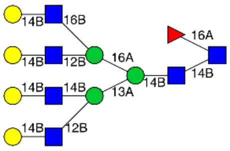 |
| T323 | 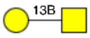 | 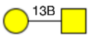 | 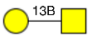 |
| S325 | 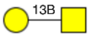 | — | — |

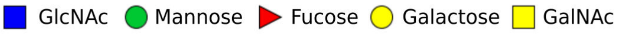

#### 2 Supplementary Figures

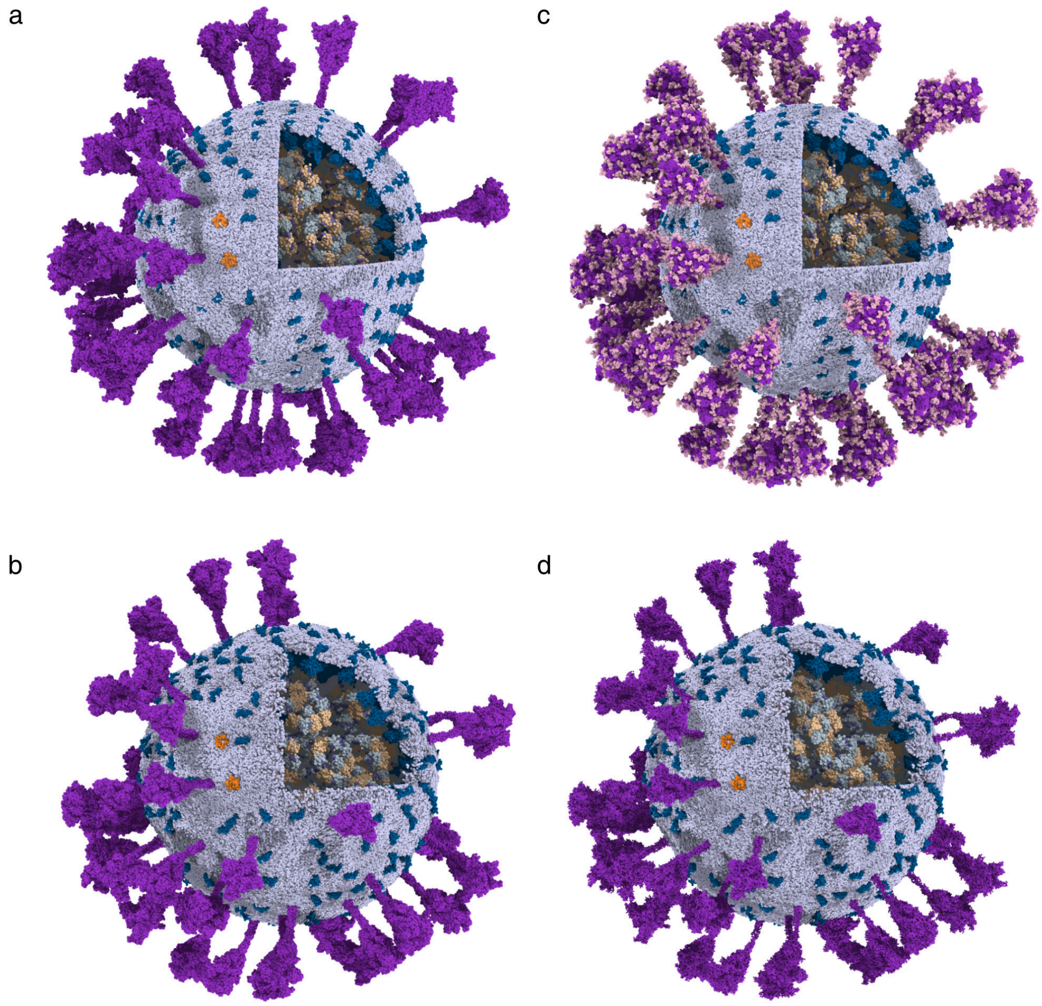


**Figure S1:** Overview of the coarse-grained and atomistic SARS-CoV-2. The viral structural proteins were colored in purple (S), deep-blue (M), orange (E), grey-blue (RBD of N), and wheat (dimerization domain of N). The viral envelope was rendered in blue-white. RNA segments were in blue-purple. In the atomistic system, the glycans were colored in light pink. (a) The initial coarse-grained structure of the virus; (b) The final coarse-grained virus structure after the 500-ns CG MD simulation; (c) The atomistic virus structure transformed from (a); (d) The atomistic virus structure transformed from (b), the S proteins were rendered in surface representation.

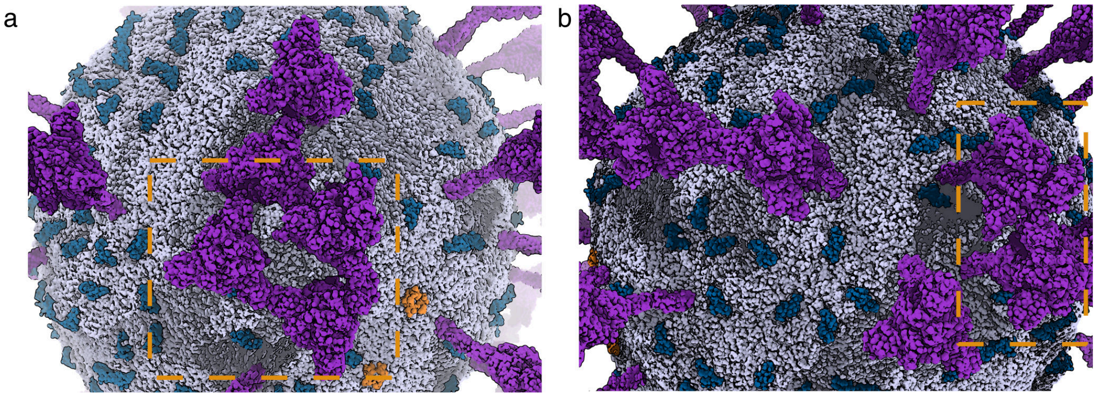


**Figure S2:** Overview of the S protein head region cluster. The S, M, E proteins were colored in purple, blue, orange, respectively. The viral envelope were painted in grey. The orange dashed boxes point out the areas that several S protein head regions contact with each others.

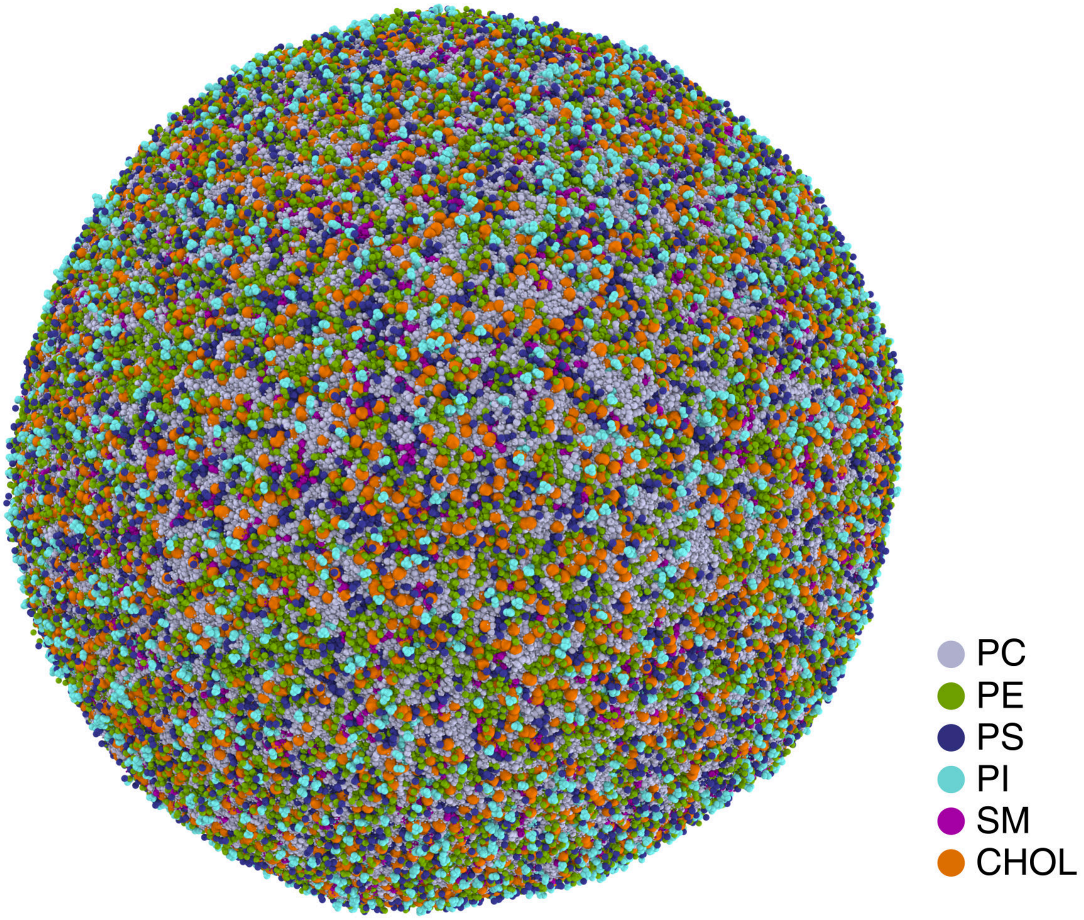


**Figure S3:** The overview of the equilibrated vesicle.

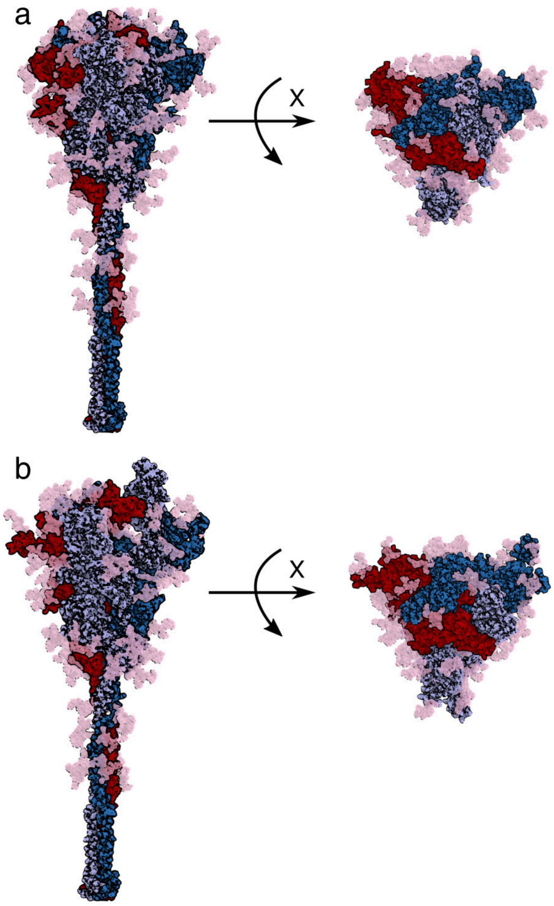


**Figure S4:** Structures of the spike protein. (a) Full-length 'RBD down' state. The three chains were colored blue-white, sky blue, and red, respectively. The glycans were colored in light pink. (b) same as (a) but for the 'one RBD up' state.

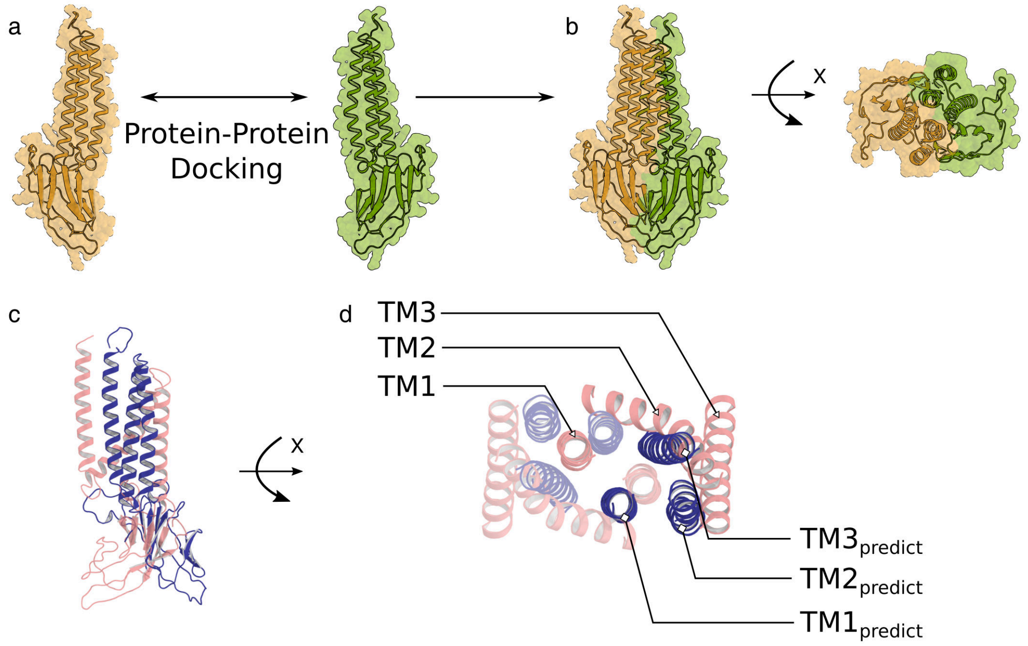


**Figure S5:** (a) The M protein monomer structure (green and orange) predicted by AlphaFold2; (b) The side and top views of the M dimer structure generated by Protein–Protein docking (ZDOCK). (c) The side view of the two M protein monomer structures: structure from docking (deepblue); 'long' structure (pink, PDB ID: 7vgr). (d) The top view of these two M dimer structures transmembrane helices.

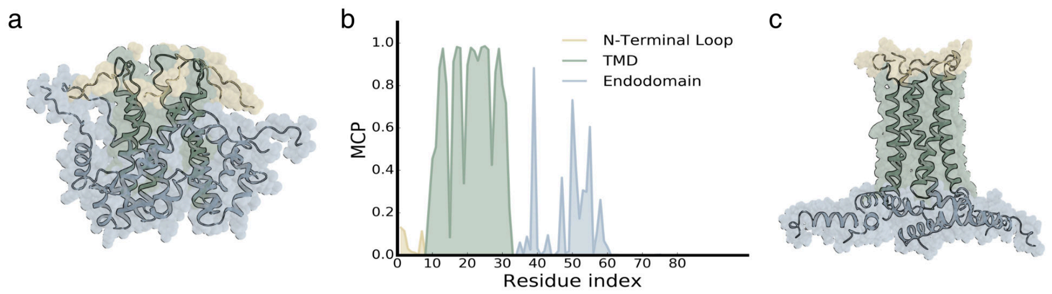


**Figure S6:** The structures of the E protein pentamer. (a) The homology modeling structure based on the SARS-CoV E protein; (b) The predicted membrane contact probability (MCP) of the E protein. (c) The E protein pentameric structure optimized by Feig's Lab. The brown, green, and blue regions represent the N-terminal loop, the transmembrane domain, and endodomain of the E protein, respectively.

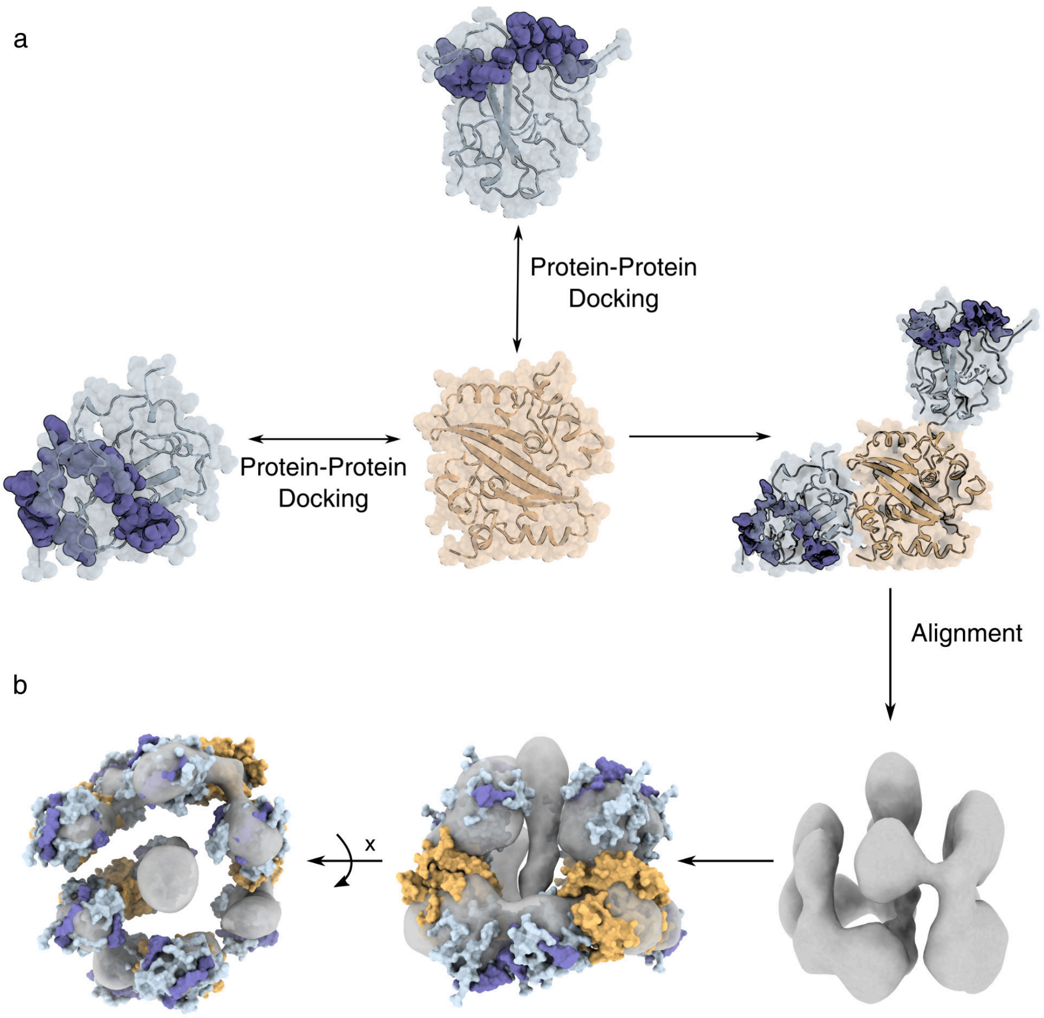


**Figure S7:** Construction of the RNP unit structure. (a) The N protein dimer (right panel) was generated by docking two NTD structures (light blue) to the dimerized CTD structure (light orange). The RNA segments bound to N proteins were colored deep and opaque blue; (b) Five N protein dimers are fitted into the Cryo-ET density (grey surface) to form a single RNP unit (left panel).

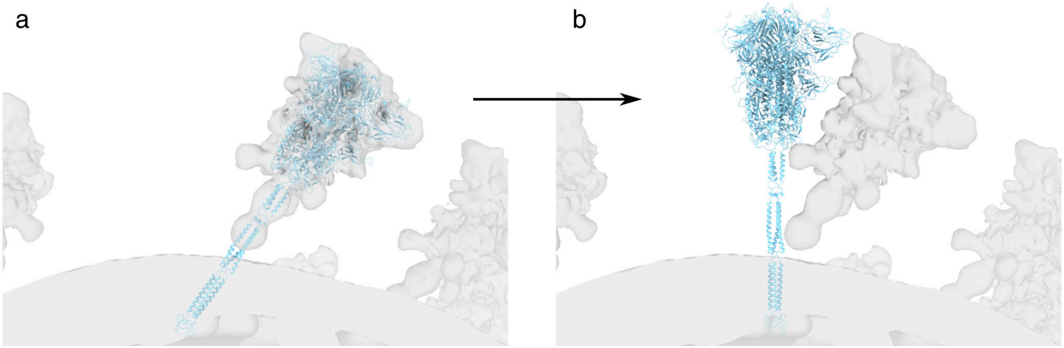


**Figure S8:** (a) Aligning the S protein into the Cryo-ET density (Chimera –> Fit into density) may lead to inappropriate positioning of the transmembrane domain of the S protein. (b) The adjusted S protein orientation with the transmembrane domain properly embedded into the lipid bilayer. The light blue cartoon indicates the structure of the S protein. The grey surface shows the Cryo-ET density.

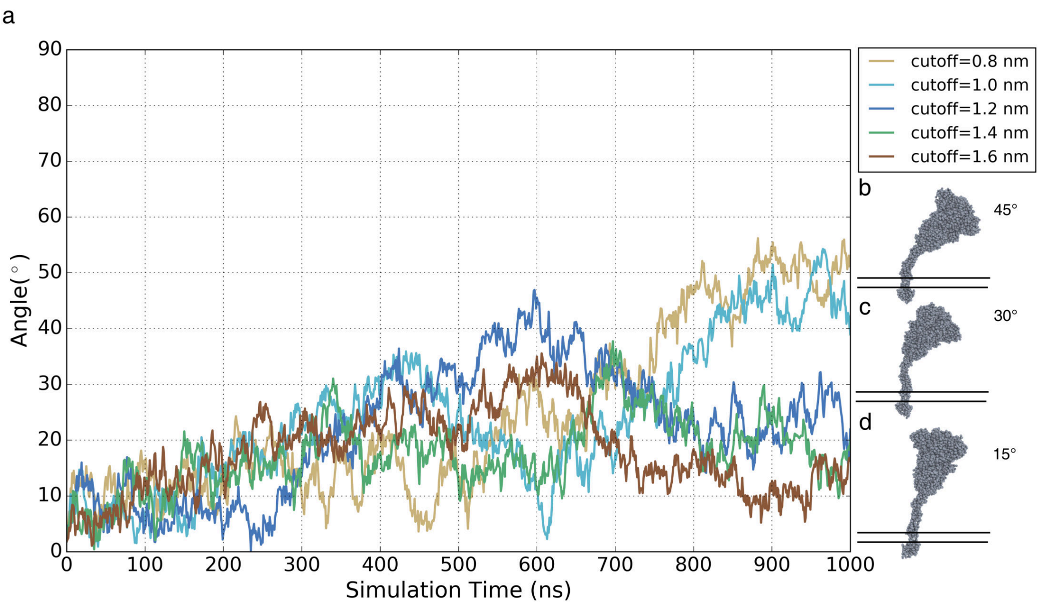


**Figure S9:** The orientation of the spike protein on the lipid bilayer depends on the elastic network cutoff. (a) Each curve elucidates the spike protein orientation distribution in CG MD simulations with different elastic network cutoff values. The angle was measured between the axis of the spike protein and the normal axis of the membrane surface. (b)–(d) The typical S protein orientations during the simulations.

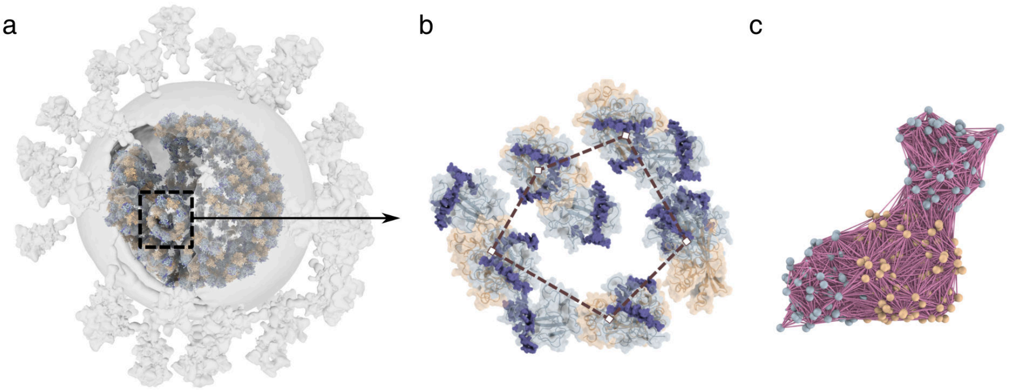


**Figure S10:** The RNPs and the distance restraints within them. (a) Alignment of the RNP units into the Cryo-ET density map. (b) The zoom-in view of the black frame in (a). The dashed lines represent the distance restraints applied to the N protein dimers in each RNP unit. (c) Magenta lines indicate the pairwise elastic network to maintain the N protein dimer conformation. Light blue and light orange represent domains near the NTD and CTD of the N proteins, respectively. Deep blue indicates the RNA segments bound to N proteins.
